# Differential effects of diet and weight on taste responses in diet-induced obese mice

**DOI:** 10.1101/564211

**Authors:** Zachary C. Ahart, Laura E. Martin, Bailey R. Kemp, Debarghya Dutta Banik, Stefan G.E. Roberts, Ann-Marie Torregrossa, Kathryn F Medler

## Abstract

The ever growing obesity epidemic has created a need to develop a better understanding of the underlying mechanisms responsible for this condition. Appetite and consumption are directly influenced by the taste system which determines if potential food items will be ingested or rejected. While previous studies have reported that obese individuals have reduced taste perception, the relationship between these processes is still poorly understood. Earlier work has demonstrated that diet-induced obesity (DIO) directly impairs taste responses, particularly for sweet stimuli. These deficits occurred in the cells located in the oral cavity as well as in the behavioral responses. However, it is not clear if these changes to the taste system are due to obesity or to the high fat diet exposure. The goal of the current study was to determine if diet or excess weight is responsible for the DIO induced taste deficits. Using a combination of live cell imaging, brief-access licking, immunohistochemistry and real-time PCR, we have found that diet and weight gain can each selectivity affect taste. Follow up experiments determined that two key signaling proteins, gustducin and phospholipase Cβ2, are significantly reduced in the high fat diet without weight gain and obese mice, identifying a potential mechanism for the reduced taste responsiveness to some stimuli. Our data indicate that the relationship between obesity and taste is complex and reveal that for some stimuli, diet alone can cause taste deficits, even without the onset of obesity.

---

The obesity epidemic and its health related complications have made it imperative that we gain a better understanding of the factors that contribute to this condition. Several studies have suggested that perception of taste stimuli is reduced in obese populations [1-8] and that decreased sensitivity to taste stimuli and food associated signals lead to increases in intake [9-11]. Despite this link between taste and obesity, little work has focused on identifying the mechanisms that are responsible for this relationship. While some studies have described the effects of obesity on the central taste system, understanding the role of peripheral taste in obesity is essential because the taste receptor cells are the site of interaction between the organism and the potential food item. Our lab previously reported that diet-induced obesity (DIO) significantly inhibits the responsiveness of peripheral taste cells to taste stimuli, particularly sweet stimuli, and that these changes translate into behavioral effects [12]. There is strong support for the idea that a reduction in taste signaling drives increases in food intake [9, 10, 13] and to our knowledge, our earlier study [12] was the first to demonstrate that DIO affects multiple aspects of taste from the initial signaling event to taste-driven behaviors.

There are two aspects of DIO that can potentially affect taste responses: excess weight or diet. We have now explored the respective roles of these two factors for the DIO-dependent taste deficits we previously identified. We focused our studies on sweet taste because it was the most affected by DIO [12] and is reported in other studies to be affected by obesity [9-11, 14-16]. The goal of this current study was to identify if there are independent effects of diet or excess weight on taste function.

In order to separate weight gain from exposure to a high fat diet, we added a low dose of captopril to the water of mice on a high fat diet. Previous work has shown that captopril (CAP), an angiotensin converting-enzyme inhibitor protects against the development of diet-induced obesity [17]. Animals on CAP voluntarily reduce their caloric consumption, which prevents weight gain. Similarly, exposing DIO animals to CAP also causes weight loss and the reversal of the associated metabolic issues that normally occur with obesity, including improved glucose tolerance and insulin resistance, loss of adipose tissue, and reduced inflammation [17-19]. This pharmacological approach to expose mice to a HF diet without weight gain avoids any potential stress that using food restriction and single housing might impose on the mouse. Importantly, our control experiments found that the concentration of CAP used in these experiments did not independently affect taste responses.

Using this approach, we compared DIO mice to mice that were on the same HF diet but did not become obese. We found that excess weight had the largest influence on taste deficits but that diet alone also impaired taste responses for some, but not all, stimuli. These data suggest a complex relationship between DIO and taste that is influenced by both weight and diet.

## Results

### High fat diet causes significant weight gain in the absence of captopril

Weekly tracking of total body weight identified a rapid weight gain in mice fed a high-fat (HF) diet. After one week, mice on HF food are significantly heavier than mice on standard chow (CTL). Mice on HF food with captopril in their water (HF+CAP), and mice on standard chow with captopril (CTL+CAP) did not rapidly gain weight and were not significantly different from CTLs (Figure 1A).

**Figure 1:**
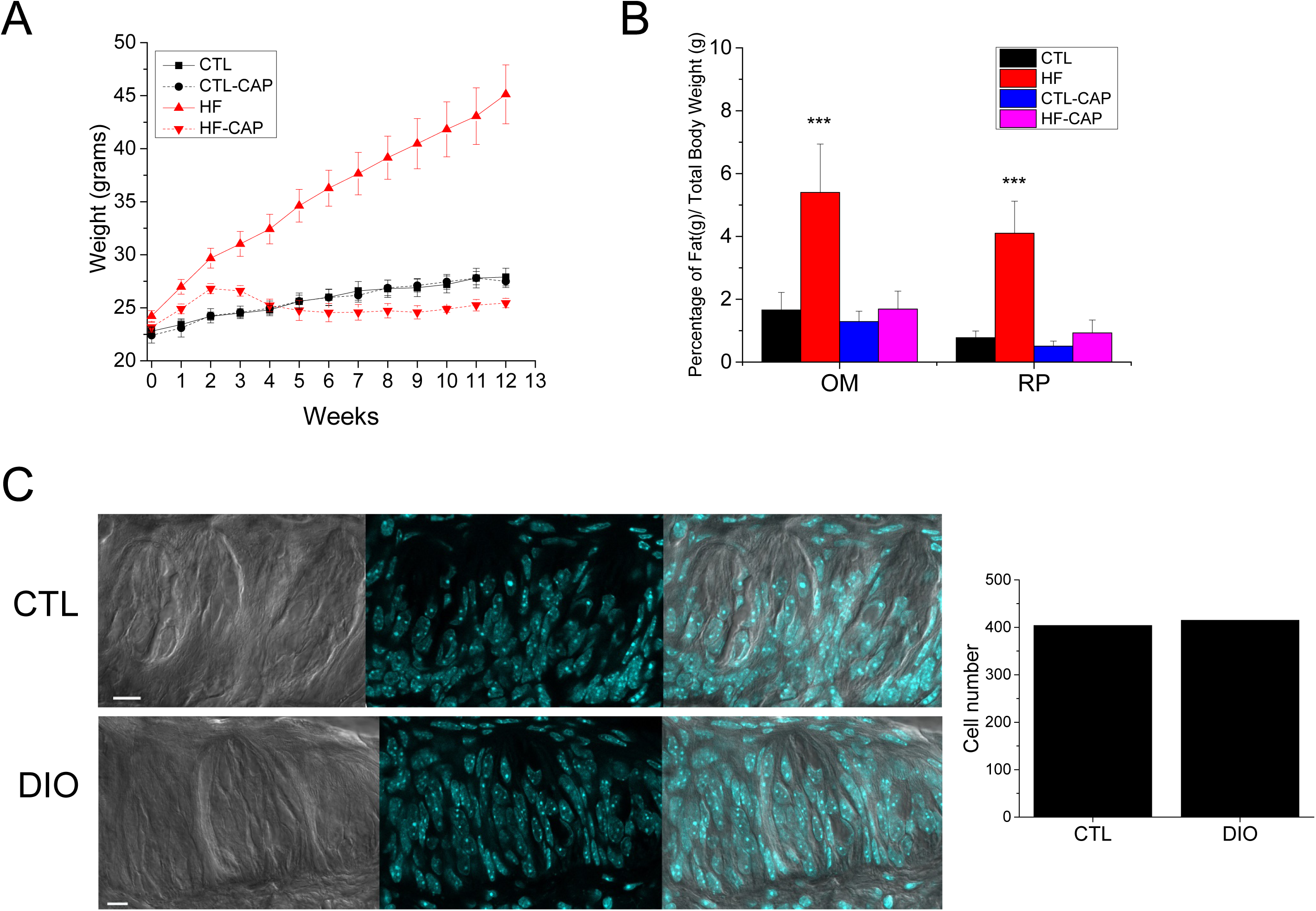
Mice on a HF diet become obese but do not gain excessive weight in they are exposed to captopril. **A**) Diet induced obesity (DIO) was defined by measuring total body weight and adipose tissue weight for mice on HF (+/−CAP) compared to CTLs. There was a significant increase in the total body weight of the HF-CAP mice by 1 week on the HF diet (p=0.01) and their weights continued to increase over time. No other significant differences were found for the other groups (n=4 for each group). **B**) The HF-CAP mice (red bar, n=10) had significantly more omentum and retroperitoneal fat as a ratio of total body weight compared to controls (***p<0.001) while mice on the HF diet with CAP (pink bar, n=10) were not different from either control (black or blue bar, n=5 for each). Data are presented as a percentage of total weight. **C**) The total number of taste cells in control and DIO mice were counted using DAPI staining to label individual taste cells (examples shown in left panels. The number of taste cells is not reduced in DIO mice (13 buds from 3 mice for each, graph on the right). DIO mice on HF diet ∼10 weeks with OM>3 and RP> 3 vs CTLs.

We assessed the level of obesity in the mice for each experimental condition by measuring the mass of the greater omentum (OM) and retroperitoneal (RP) adipose tissues (Figure 1B). These measurements are reported as a percentage of total mouse weight and identified a significant increase for both the ratio of OM/total body weight (one way ANOVA, p<0.001) and the RP/total body weight in mice on the HF diet alone compared to mice in the other experimental conditions (one way ANOVA, p<0.001). No other significant differences were found. Based on these criteria (overall weight and mass of these adipose tissues), mice on the HF (-CAP) diet were identified as obese.

### Diet induced obesity does not cause loss of taste receptor cells

To determine if diet-induced obesity (DIO) caused a significant loss of the peripheral taste cells, control experiments analyzed taste buds from mice (n=3) that were on the HF diet for 10 weeks and had OM>3, RP>3. The number of taste cells per bud in these mice were compared to age matched littermates on CTL diet (n=3). Taste buds from the circumvallate papillae (CV) were fixed and stained with DAPI to identify the nuclei (Figure 1C) for cell counting. 40 micron sections were imaged at every micron and every 5^th^ section was counted. Cell counting was performed with the experimenter blind to condition. A total of 13 taste buds were analyzed for each condition and no significant differences were found between the CTL or DIO mice (Figure 1C, bar graph).

### CAP alone does not significantly alter taste driven behaviors

Preliminary studies were used to ensure that CAP alone was not altering taste driven behaviors. We performed control experiments with mice that were pre-exposed to CAP for 4 days while on standard chow before they were behaviorally tested. There were no differences in the taste-driven behaviors in the unconditioned licking experiments to sucrose in these mice compared to mice that were not exposed to CAP (Supplemental Figure 1). We concluded that exposure to CAP alone does not significantly affect taste driven behaviors.

### Diet and weight can have differential effects on sweet taste responses

We then evaluated the effects of excess weight and exposure to HF diet for three sweet stimuli, two artificial sweeteners: acesulfame K (AceK, 20mM) and saccharin (sac, 2mM) as well as sucrose (suc, 50mM). A bitter compound, denatonium benzoate (5mM), and 50mM KCL were also tested. Three different measures were taken to test the effects of obesity and diet on taste: 1) the percentage of taste cells that respond to a given stimulus, 2) the amplitude of the taste evoked calcium signal in the cells that responded to the stimulus, and 3) the unconditioned licking response in the behaving animal for each stimuli (Figures 2 and 3).

**Figure 2:**
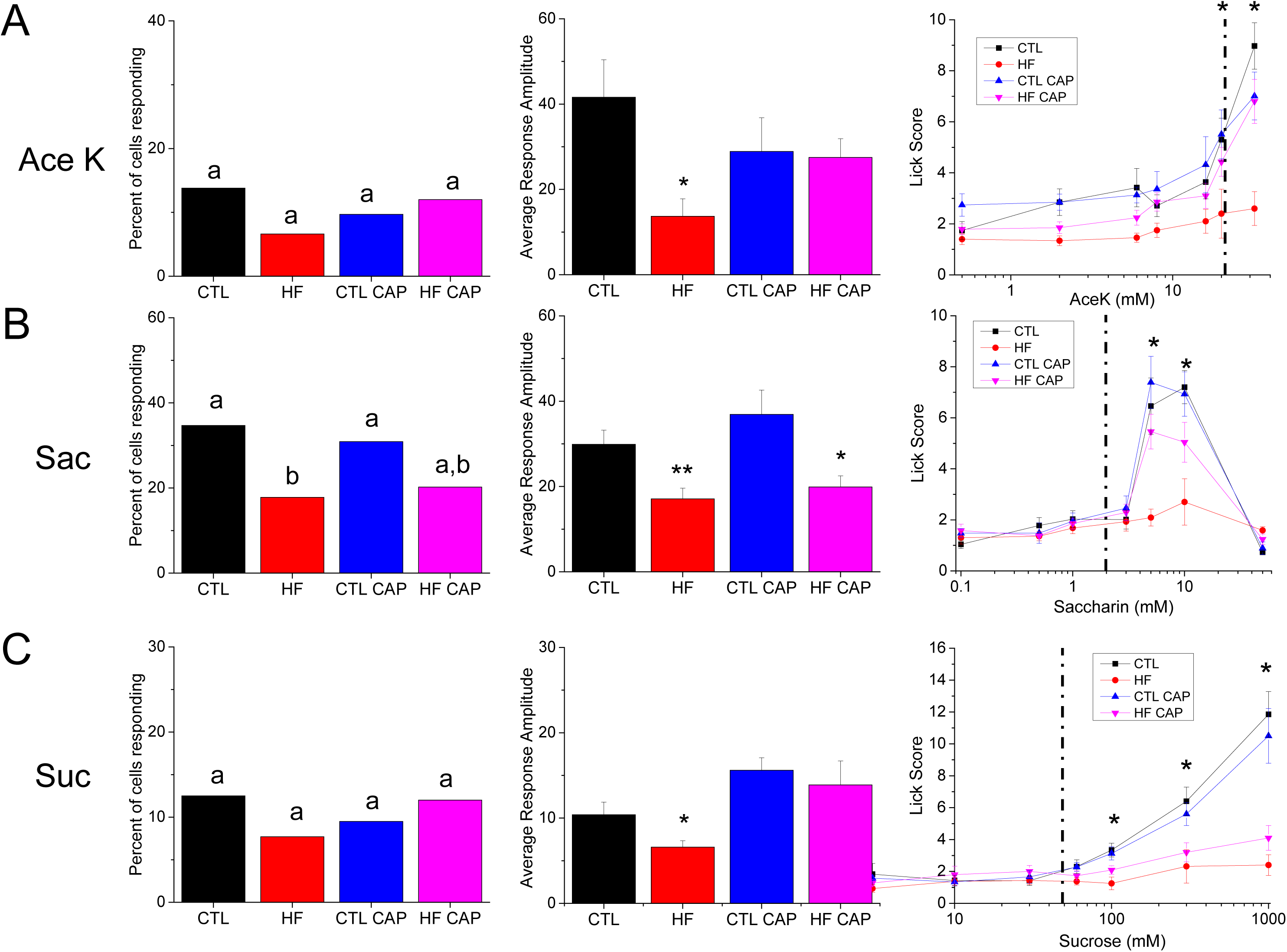
Differential effects of diet and weight on sweet stimuli. A combination of live cell imaging and unconditioned brief-access licking assays were used to assess the relative effects of DIO and HF diet on the ability of the taste system to respond to sweet stimuli. The left column is the total number of taste cells that responded to each stimulus. HF diet (+/−CAP) were compared to CTL (+/−CAP) mice using Chi-square analysis with Yates correction (significant differences are indicated by different letters). The center column is the response amplitude for each condition (compared with one-way ANOVA) and the third column is brief-access licking behavior for each stimulus (compared with ANOVAs, see Table 1). The dotted line in the third column indicates the stimulus concentration used in the live cell imaging experiments. Two artificial sweeteners, Ace K (20mM, **A**) and saccharin (sac, 2mM, **B**) were evaluated along with sucrose (suc, 50mM, **C**). Individual data values for each condition is shown in Supplemental Table 1. **A**) No significant differences were found in the number of taste cells that responded to AceK, however response amplitudes from the HF mice were significantly lower than CTL (p=0.012) and HF+CAP (p=0.03). Consistent with the amplitude data, HF+CAP mice licked significantly less than the other treatments in the brief-access test. **B**) Treatment with a HF diet reduced the percent of cells responding to saccharin (p=0.04), as well as the average response amplitude (one way ANOVA, p=0.001). In the brief-access taste test HF mice licked significantly less than other groups and the HF+CAP group trended to lower licking behavior (p=0.053) compared to the CTL animals. **C**) There was no effect of weight or diet on the percentage of cells responding to sucrose, while there was a reduction in response amplitude in HF animals (one way ANOVA, p=0.02). The behavioral data suggest that exposure to the HF diet is sufficient to alter licking behavior.

**Figure 3:**
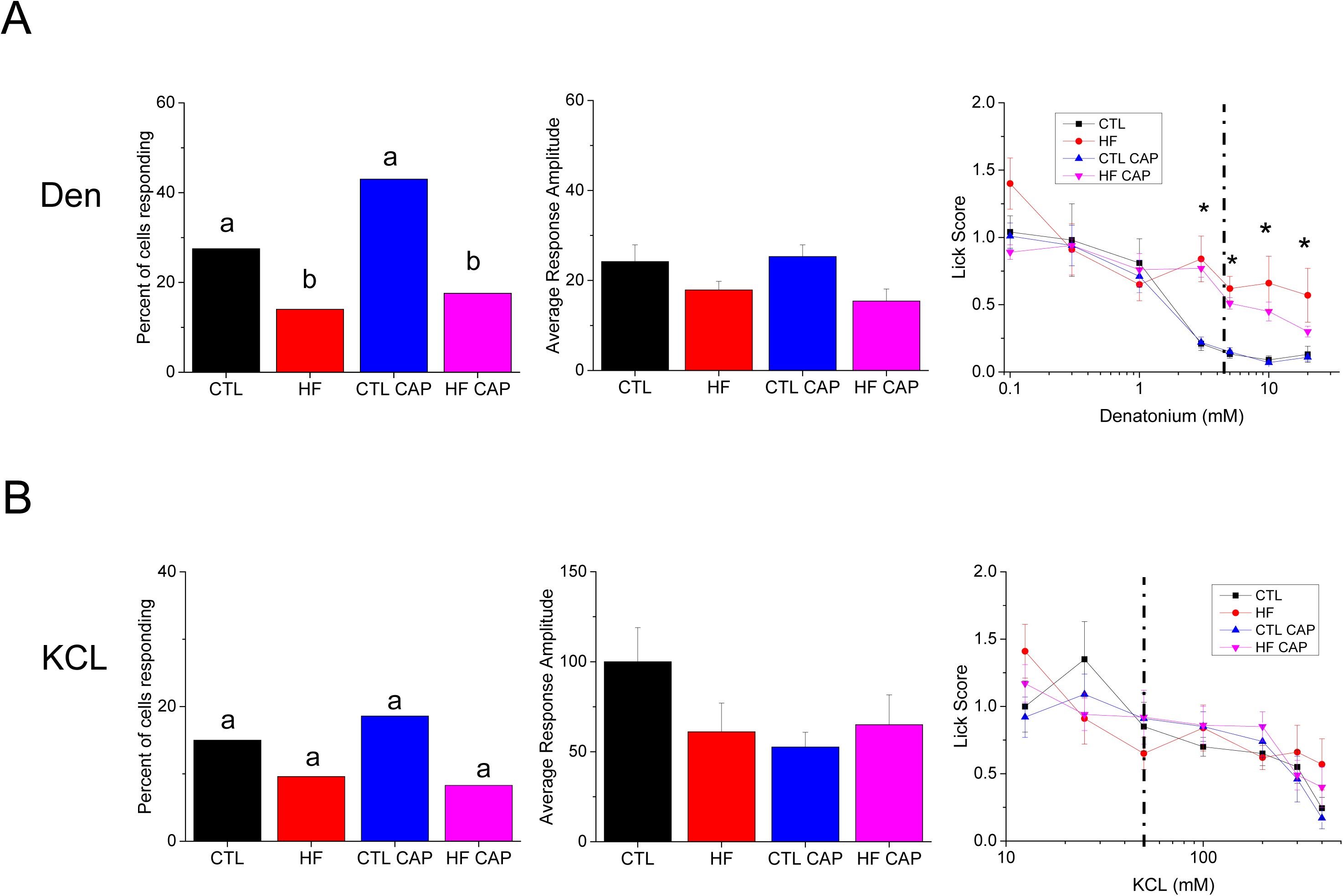
Effects of diet and weight on bitter and salt stimuli. A combination of live cell imaging and unconditioned brief-access licking assays were used to assess the relative effects of DIO and HF diet on the ability of the taste system to respond to bitter (5mM Denatonium Benzoate, **A**) and salt (50mM KCL, **B**) stimuli. Individual data values for each stimulus is shown in Supplemental Table 1. The left column is the total number of taste cells that responded to each stimulus. Mice on the HF diet (+/−CAP) were compared to CTL (+/−CAP) mice using Chi-square analysis with Yates correction). The center column is the response amplitude for each condition (compared with one-way ANOVA) and the third column is brief-access licking behavior for each stimulus (compared with ANOVAs, Table 1). The dotted line in the third column indicates the stimulus concentration used in the live cell imaging experiments. **A**) Exposure to the HF diet reduced the percentage of cells responding to Den (p=0.029 for HF-CAP and p=0.007 for HF+CAP), and decreased the behavioral sensitivity to Den, but did not affect the response amplitudes of the responses. **B**) There was no effect of diet or weight on any measure of responding to KCL.

**Table 1.** Statistical data for the unconditioned licking behavioral data shown in Figures 2 and 3.

| Hypothesis Group |  | all groups comparison<br>all groups |  |  | effect of CAP in LF<br>CTL ± CAP |  |  | effect of HF diet<br>CTL vs HF+CAP |  |  | effect of weight<br>CTL vs HF-CAP |  |  | effect of cap in HF<br>CTL ± CAP |  |  |
| --- | --- | --- | --- | --- | --- | --- | --- | --- | --- | --- | --- | --- | --- | --- | --- | --- |
|  | effect | F | df | p | F | df | p | F | df | p | F | df | p | F | df | p |
| **ace k** | treatment | 15.613 | (3,28) | <0.001 | 0.004 | (1,14) | 0.947 | 2.777 | (1,22) | 0.11 | 19.14 | (1,22) | <0.001 |  |  |  |
|  | concentration | 23.848 | (6,168) | <0.001 | 25.786 | (6,84) | <0.001 | 33.83 | (6,132) | <0.001 | 32.449 | (6,132) | <0.001 |  |  |  |
|  | interaction | 6.928 | (18,168) | <0.001 | 1.587 | (6,84) | 0.161 | 0.36 | (6,132) | 0.903 | 5.873 | (6,132) | <0.001 |  |  |  |
| **saccharin** | treatment | 5.663 | (3,28) | 0.004 | 0.214 | (1,14) | 0.651 | 1.498 | (1,22) | 0.234 | 15.968 | (1,22) | 0.001 |  |  |  |
|  | concentration | 81.404 | (6,168) | <0.001 | 55.981 | (6,84) | <0.001 | 62.198 | (6,132) | <0.001 | 38.013 | (6,132) | <0.001 |  |  |  |
|  | interaction | 5.244 | (18,168) | <0.001 | 0.368 | (6,84) | 0.898 | 2.142 | (6,132) | 0.053 | 14.854 | (6,132) | <0.001 |  |  |  |
| **sucrose** | treatment | 20.71 | (3,28) | <0.001 | 1.587 | (1,14) | 0.228 | 22.587 | (1,22) | <0.001 | 47.005 | (1,22) | <0.001 | 4.622 | (1,14) | 0.50 |
|  | concentration | 73.212 | (6,168) | <0.001 | 86.372 | (6,84) | <0.001 | 49.475 | (6,132) | <0.001 | 40.186 | (6,132) | <0.001 | 5.24 | (6,84) | <0.001 |
|  | interaction | 12.452 | (18,168) | <0.001 | 1.287 | (6,84) | 0.272 | 18.742 | (6,132) | <0.001 | 24.715 | (6,132) | <0.001 | 0.5 | (6,84) | 0.806 |
| **DB** | treatment | 9.922 | (3,28) | <0.001 | 0.17 | (1,14) | 0.686 | 12.574 | (1,22) | 0.002 | 27.208 | (1,22) | <0.001 | 3.525 | (1,14) | 0.081 |
|  | concentration | 39.811 | (6,168) | <0.001 | 28.4 | (6,84) | <0.001 | 30.064 | (6,132) | <0.001 | 20.973 | (6,132) | <0.001 | 13.649 | (6,84) | <0.001 |
|  | interaction | 2.516 | (18,168) | 0.001 | 0.055 | (6,84) | 0.999 | 3.918 | (6,132) | 0.001 | 5.068 | (6,132) | <0.001 | 0.86 | (6,84) | 0.528 |
| **KCI** | treatment | 0.144 | (3,28) | 0.933 | 0.048 | (1,14) | 0.830 |  |  |  |  |  |  |  |  |  |
|  | concentration | 20.648 | (6,168) | <0.001 | 13.587 | (6,84) | <0.001 |  |  |  |  |  |  |  |  |  |
|  | interaction | 1.529 | (18,168) | 0.108 | 0.609 | (6,84) | 0.723 |  |  |  |  |  |  |  |  |  |

Taste cell responsiveness is the number of taste cells that respond to a specific stimulus divided by the total number of taste cells that were exposed to that stimulus. These data are presented as a percentage and are shown in the first column of Figure 2 for Ace K (Figure 2A), saccharin (Figure 2B) and sucrose (Figure 2C). Individual values are shown in Supplemental Table 1. Chi-square analyses [20] were used to identify any statistical differences in the taste cell responsiveness between the CTL mice (+/−CAP) and the mice on HF food (+/−CAP). No significant differences were found in the number of taste cells that responded to AceK (Figure 2A). However, the average amplitude of the taste cell responses to AceK was significantly reduced in the mice on the HF diet (-CAP) compared to the other conditions (Figure 2A, middle panel, one way ANOVA, p=0.03). No other significant differences were found. Unconditioned licking was significantly reduced in the obese animals (HF-CAP) compared to the lean animals (Table 1). These data indicate that obesity is likely the primary factor resulting in the reduced taste response to the artificial sweetener AceK.

Analyses of the taste responses to the artificial sweetener saccharin found that the number of taste cells that responded to saccharin was significantly reduced in the obese mice compared to control (Figure 2B, left panel, p=0.04); however, it was not significantly different from number of saccharin responsive cells in the HF+CAP mice. While the number of taste cells from the HF+CAP mice that were responsive to saccharin was lower than the CTL-/+CAP mice, there was no significant differences identified. Evaluation of the response amplitudes identified significant differences (Figure 2B, middle panel, one way ANOVA, p=0.001). Follow up Student’s t tests determined that the saccharin responses in the obese mice were significantly smaller than CTL (p=0.003) with comparable reductions in the HF+CAP mice compared to the CTL+CAP mice (p=0.017). No significant differences between the mice on the HF diet (+/− CAP) or between the CTL (+/− CAP) groups were found. Unconditioned licking was significantly reduced in the obese mice (HF-CAP) compared to the lean and there was strong trend (p=0.053) for a reduction in responding in the HF+CAP mice compared to lean. Unlike, the AceK results, the saccharin data suggests that diet is driving the reduction in saccharin responses since mice that were placed on the HF (+CAP) had impaired saccharin responses comparable to the obese mice. Their behavioral responses suggest that the diet alone has an effect on the taste driven behavior but that this effect is magnified when the animal is also obese.

We repeated the same analyses for sucrose and found no differences in the percentage of responsive taste cells while the amplitudes of the sucrose evoked responses were only reduced in the obese mice. Interestingly, in the behavioral analysis, both animals on the HF diet (+/−CAP) showed reduced unconditioned licking compared to the CTL mice (+/− CAP) as sucrose concentration increases. This suggests that the decrease in licking is due to the exposure to the HF diet, not changes in body mass.

### Diet affects bitter taste but not KCL taste

We also tested the bitter compound denatonium (5mM) and 50mM KCL which is perceived as salty. Exposure to HF diet regardless of weight change, significantly reduced the number of taste cells that responded to denatonium (Figure 3A, left panel) but did not have a significant effect on the amplitude of the denatonium evoked signals (Figure 3A, middle panel). The behavior data demonstrates that exposure to the HF diet reduced their sensitivity (increased unconditioned licking) to denatonium (Figure 3A, right panel and Table 1). In agreement with our earlier study [12], 50mM KCL responses were not affected by weight gain or diet exposure for any of the parameters analyzed (Figure 3B).

### Taste signaling proteins change in obese mice

To investigate the potential reasons that the taste evoked calcium responses were inhibited, we measured the expression of α-gustducin and phospholipase Cβ2 (PLCβ2), two proteins in the signaling pathway that transmits sweet and bitter taste signals within taste receptor cells. Immunohistochemical analysis suggested that there was a reduction in the expression of both of these signaling proteins, so further experiments quantitated the number of taste cells expressing α-gustducin and PLCβ2 in the CV taste buds for mice from each experimental condition. Analysis parameters were standardized to DAPI labeling which was quantified as described above and was done with the experimenter blind to condition. After the total number of taste cells/bud were determined, the number of cells with labeling for each protein of interest was measured (n=at least 3 mice for each). Gustducin and PLCβ2 were analyzed separately.

The relative expression of gustducin for each experimental condition is shown in Figure 4A, left panel. Chi-square analysis identified a significant reduction in the number of cells expressing α-gustducin in obese taste buds (HF) compared to controls (p=0.02). This indicates that there are fewer cells expressing gustducin in obese mice (Figure 4A, middle panel). No other significant differences were found. Real-time PCR was used to quantitate the mRNA levels for gustducin from isolated taste buds (n=3 mice for each experimental condition). There was a significant reduction in gustducin transcript levels in both the HF +/−CAP mice (Figure 4A, right panel, **, p<0.01) compared to CTLs.

**Figure 4:**
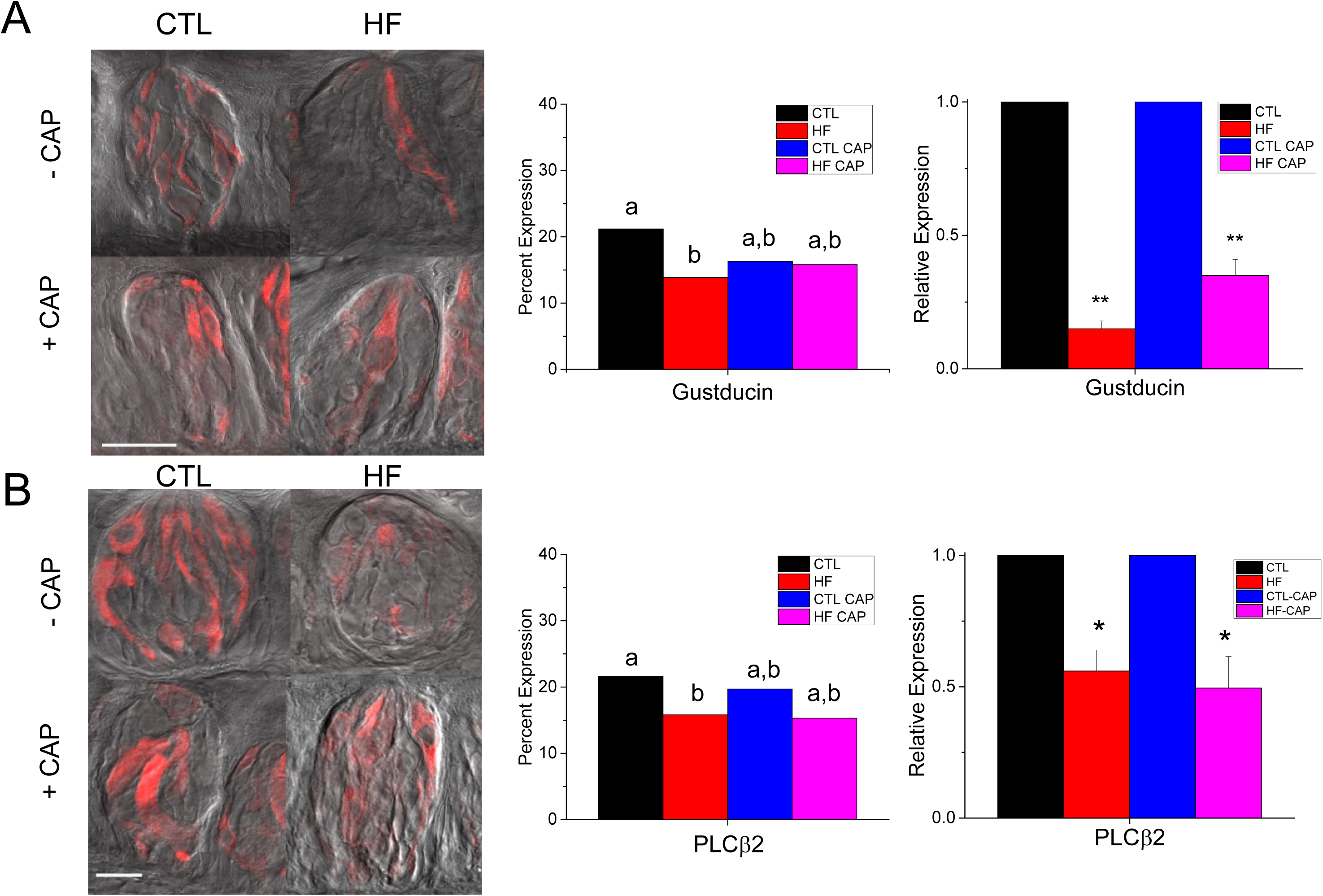
α-Gustducin and PLCβ2 expression are affected by DIO. **A**) Representative immunohistochemical images of α-gustducin expression (red) in the CV taste buds of HF+/− CAP and CTL+/− mice, as well as HF-CAP and CTL (left panel, scale bar=20μm). Quantitative analysis determined that there is a significant decrease in the number of α-gustducin-expressing cells in obese mice compared to controls (middle panel, p=0.02). No other significant differences were found. qPCR analysis identified a significant reduction in gustducin mRNA levels in both HF (**, p=001) and HF+CAP mice (**, p=0.009). **B**) Representative immunohistochemical images of PLCβ2 in CV sections of control and obese mice +/−CAP (left panel, scale bar=20μm). There is a significant reduction in the expression of PLCβ2 in cells from obese mice compared to CTL mice (middle panel, p=0.04). No other significant differences were found. There is a significant reduction in the mRNA expression of PLCβ2 in obese taste cells compared to controls (right panel, *, p=0.035) as well as the mice on the HF +CAP (right panel, *, p=0.013).

Immunohistochemical data in Figure 4B (left panel) suggests that the PLCβ2 expression is reduced for the CV taste buds from obese mice. Analysis of the PLCβ2 expression identified a significant reduction in the number of CV taste cells expressing PLCβ2 in the obese mice compared to CTL (Figure 4B, middle panel, p=0.04). Even though the number of PLCβ2 expressing cells in the HF CAP mice was reduced, it was not significantly different from CTLs. Real-time PCR was used to measure the mRNA levels for PLCβ2 from isolated taste buds (n=3 mice for each experimental condition). PLCβ2 transcript levels were significantly reduced in the HF +/−CAP mice (Figure 4B, right panel, *, p<0.05) compared to CTLs.

## Discussion

Obesity is a complex disease and its relationship with the taste system is still not well understood. While multiple studies have suggested that perception of taste stimuli is reduced in obese populations [1-8] and that this leads to increased consumption [9-11], the mechanisms underlying these effects have not been identified. We have now demonstrated that a HF diet and excess weight can each inhibit taste responses and that these effects vary depending on the stimulus. Other studies have suggested a role for both leptin resistance and low grade inflammation in obesity-related changes in taste sensitivity [21, 22]. While this likely true for some taste stimuli, our data demonstrates that a prolonged exposure to a high fat diet, even in the absence of obesity, can reduce the sensitivity of the taste system to certain stimuli.

We measured the effects of DIO and diet on both the activity of taste receptor cells, which is the first step of the taste pathway, and the behavioral responses, which are the final output of the taste pathway. For each stimulus, we ensured that the behavioral experiments tested a range of concentrations that encompassed the stimulus concentration that was used in the live cell imaging experiments (shown as a dotted line on the behavior graphs). For some stimuli (AceK and denatonium), the largest separation in the behavior data correlated with the concentrations used in the imaging experiments. However, for other stimuli (saccharin and sucrose), the concentration used in the imaging experiments was lower than the stimulus concentrations where behavioral differences were recorded. While we used these concentrations to avoid non-specific effects that can occur in live cell imaging, it is possible there would be more significant differences in the cell responses if a higher stimulus concentration could be used. Indeed, the behavioral results for sucrose indicate that diet alone causes a comparable impairment to the obese mice at higher concentrations (>100mM) even though there was not a measureable difference in our live cell imaging experiments (50mM).

While high concentrations of saccharin taste bitter, we do not believe that our findings reflect a bitter specific effect since the saccharin concentration we used is well below the threshold that is perceived as bitter [23-25] as our behavioral data confirms. We also recorded a diet effect on sucrose behavioral responses that further demonstrates the diet effect is not bitter specific. The strong agreement between our cellular and behavioral data supports the premise that the diet dependent inhibition of the taste cell activity is significant enough to impair the animal’s behavior. Thus, diet alone can alter taste driven behaviors, even in the absence of weight gain. This occurs, at least in part, by altering the properties of the taste receptor cells.

Interestingly, our results for denatonium, the bitter compound that we tested in this study, had some differences from our previous findings [12]. In our first study, there was a reduction in the number of denatonium responsive cells in the obese mice, but it was not significant differently from CTL. However, in our current study, obese mice did have a significant reduction in the percentage of cells that responded to denatonium compared to CTL (Figure 3A). Both studies identified significant differences in the denatonium driven behaviors. These differences in cell responsiveness are likely due to differences in the number of taste cells evaluated in the different studies since more cells were tested in the current study. Obesity reduced the behavioral responses in both studies and we have now shown that the loss of aversion to denatonium is at least in part, influenced by diet.

To begin identifying the potential cellular mechanisms underlying the changes in taste cell activity, we evaluated the expression levels of two proteins with known roles in the transduction of bitter and sweet stimuli. α-Gustducin is a G-protein that is expressed in 20-30% of taste cells and is required for the normal transduction of these stimuli [26]. We found that the number of taste cells expressing the gustducin protein was significantly reduced in the obese mice but was not significantly reduced in the mice on the HF diet with no weight gain (Figure 4A). Conversely, qPCR for α-gustducin identified a significant reduction in mRNA in taste cells from both HF+CAP and obese mice. Since there was not a significant reduction in the number of gustducin expressing cells in the HF+CAP mice, we predict that the level of gustducin expression within the individual taste cells of these mice may be reduced. Loss of gustducin significantly decreases the ability to detect bitter and sweet stimuli [27], so the diminished gustducin expression in the obese mice is likely contributing to their impaired ability to respond to the sweet and bitter stimuli we tested. Since gustducin is not solely responsible for the transduction of all sweet stimuli [28], this selective reduction in its expression may be the basis, or at least contribute, to the selective diet effect on the sweet taste responses that we found.

We also evaluated the expression of PLCβ2. This enzyme produces IP_3_, which activates IP_3_ receptors on the endoplasmic reticulum to release intracellular calcium. This PLCβ2 function is also required for the transduction of bitter and sweet stimuli [29] and we found that its expression was significantly reduced in the obese mice at both the protein and mRNA levels (Figure 4B). While there appeared to be fewer cells expressing PLCβ2 in the HF+CAP mice, no significant differences were identified (p=0.05) even though there was a significant reduction in the mRNA levels for PLCβ2 (p=0.013). We postulate that the modest reduction of PLCβ2 expression in these taste cells may be a contributor to the selective effects of diet in inhibiting taste responses. These data also suggest that the obesity and diet dependent inhibition of the taste cell activity are not specific to one protein, but to many components of the taste pathway. While one sweet sensitive pathway is known and well-studied [30], the differential effects of the HF diet for the different sweet taste stimuli, suggests different sweet-sensitive pathways may exist. This idea is supported by earlier work in other labs [31-35].

Our study demonstrates that the peripheral taste system can be significantly inhibited by diet, without the onset of obesity. This leads to the idea that diet exposure, even without increased consumption, may be impairing taste responses sufficiently to alter consumption which can then subsequently induce obesity. Further studies are needed to address this question.

## Materials and Methods

### Mice

Animals were cared for in compliance with the University at Buffalo Institutional Animal Care and Use Committee. All experiments used C57BL/6 mice that ranged in age from 1 to 6 months. At weaning, mice were placed on either standard (Harlan labs: diet is comprised of 18% calories from fat, 58% calories from carbohydrates, 24% calories from protein) or high fat (HF) chow (60% high fat Kcal feed, Harlan Labs, Inc., Madison, WI, USA; diet is comprised of 60% calories from fat, 22% calories from carbohydrates, 18% calories from protein). Mice were placed on the HF diet (60%kcal) in the presence and absence of captopril (CAP, 0.05mg/mL water) with control mice on standard chow +/− CAP. CAP water was replaced every other day. Experiments were performed after the mice on the HF diet became obese, usually at least 6-8 weeks after they were given the HF diet. Omentum and retroperitoneal fat weights were collected and were calculated as either [(OM g/total weight g)*100] or [(RM g/total weight g)*100].

### Taste cell isolation

Taste receptor cells were harvested from taste papillae of adult mice as previously described [12, 36-41]. Briefly, mice were sacrificed using carbon dioxide and cervical dislocation. Tongues were removed and injected beneath the lingual epithelium with 0.2 mL enzyme solution (3 mg Dispase II, 0.7 mg collagenase B (Roche diagnostics, Indianapolis, IN, USA) and 1 mg Trypsin Inhibitor in 1mL Tyrode’s). Tongues were incubated at room temperature for 18 minutes in oxygenated Tyrode’s before the epithelial layer of the tongue was peeled and placed in Ca^2+^-Mg^2+^-free solution for 25-30 minutes. Cells were removed from taste papillae via gentle suction and placed on a slide coated in Cell Tak (Discovery Labware, Bedford, MA USA).

### Calcium Imaging

Isolated taste cells were incubated for 20 minutes in 2 μM Fura 2-AM (Invitrogen, Eugene, OR, USA) and nonionic dispersing agent, Pluronic F-127 (Invitrogen) and then washed for 20 minutes. Isolated cells were visualized using an Olympus IX71 scope and 40x oil immersion objective. Taste cells were stimulated and Ca^2+^ changes were recorded every 4 seconds using a Sensicam QE camera at 340/380 nm excitation, 510 nm emission. Data was collected using Imaging Workbench 6.0 (Indec Biosystems, Santa Clara, CA, USA) and analyzed using Origin 8.6 (Origin Lab Corporation, Northampton, MA, USA). Response amplitude was calculated as: [(peak value-baseline value)/baseline value]*100. Only isolated taste cells with a resting Ca^2+^ baseline between 80-150nM were analyzed and an evoked response was defined as measurable if the increase in fluorescence was greater than two standard deviations above baseline.

### Analysis of Licking Behavior

We recorded the unconditioned licking responses to varying concentrations of taste stimuli in a test chamber designed to measure brief-access licking (Davis MS80 Rig; Dilog Instruments and Systems, Tallahassee, FL). This apparatus consisted of a Plexiglas cage with a wire mesh floor. An opening at the front of the cage allowed access to one of sixteen spill-proof glass drinking tubes that reside on a sliding platform. A mechanical shutter opened and closed to allow the mouse access to one of the tubes for a user-specified length of time. A computer controlled the movement of the platform, order of tube presentation, opening and closing of the shutter, duration of tube access and interval between tube presentations. Each individual lick was detected by a contact lickometer and recorded on a computer via DavisPro collection software (Dilog Instruments and Systems).

Mice were adapted to the test chamber and trained to drink from the sipper tubes for 8 consecutive days. During training, mice were 20-h water deprived. On the first day of training, the mouse was presented with a single stationary bottle of water for 30 min. On the second day, a tube containing water was presented but this time the mouse was given 180s to initiate licking and once licking was recorded the mouse was given 30s access to the tube. At the conclusion of either the 30s access or the 180s limit, the shutter was closed again for 10s. Each of the 8 tubes, all containing water, was presented 3 times. During the remaining 5 days of training, the mouse was given 30 min to initiate licking to one of eight tubes of water. Once the mouse began licking, it was given 10s to lick before the shutter closed for 10s, after which a new tube was presented.

During testing, animals were allowed to take as many trials as possible in 30 min. Mice were tested on varying concentrations of sucrose (0,3,10,30,60,100,300,1000 mM), saccharin (0,0.1,0.5,1,3,5,10,50 mM), acesulfame K (0,0.5,2,6,8,16,20,32 mM), KCL (0,12.5,25,50,1000,200,300,400 mM), and denatonium benzoate (0,0.1,0.3,1,3,5,10,20 mM), in that order. Each stimulus was presented in randomized blocks on Monday, Wednesday and Friday in a single week. Animals were 22-h water deprived for all testing except sucrose, when animals were tested water replete. Once the animal began licking the tube, they were allowed 10 seconds of access before the shutter closed.

For stimuli tested in the water deprived condition (saccharin, acesulfame K, KCL, and denatonium benzoate), lick ratios were calculated by dividing the average number of licks at each concentration by the average number of licks to water. For stimuli tested while the animals were water replete (sucrose) licks scores were calculated by subtracting the average number of licks at each concentration by the average number of licks to water. These corrections are used to standardize for individual differences in lick rate and are based on water-need. Lick scores were compared by ANOVA with treatment (HF+CAP, HF-CAP, CTL+CAP, CTL-CAP) and solution concentration as factors. If there was a significant interaction between treatment and concentrations, these tests were followed with sub-ANOVAs. Comparisons between CTL ± CAP ruled out any potential effects of CAP. CTL treated animals were collapsed and then compared to either the HF+CAP group (to determine an effect of diet alone) or to HF-CAP (to determine the effects of weight). Bonferoni corrected t-tests were conducted to identify concentrations that differed between groups.

### Immunohistochemistry

Tongues were removed from euthanized mice and placed in 4% paraformaldehyde/0.1M Phosphate buffer (PB, pH 7.2) for 2 hours at room temperature (RT). Sucrose (20%) was then added to the fix solution and tongues were cryoprotected overnight at 4°C. The next day, 40μm sections were cut and washed in PBS 3×10 min at RT before being incubated in blocking solution (0.3% Triton X-100, 1% normal goat serum and 1% bovine serum albumin in 0.1M PBS) for 1 hour. Samples were then incubated in primary antibody for 2 hours at RT before being left overnight at 4°C. All primary antibodies were diluted in blocking solution and controls with no primary antibody were included in each experiment. The next day, sections were washed 3×10 min in PBS and incubated in secondary antibody for 2 hours in blocking solution. After 3×10 min PBS washes, sections were mounted in Flouromount media with DAPI staining (Southern Biotechnology Associates, Birmingham, AL, USA) and visualized using confocal microscopy (Zeiss Axioimager Z1 and Axiovert 200M). Primary antibodies tested: α-gustducin (1:50, Santa Cruz Biotechnology, Santa Cruz, CA, USA) and PLCβ2 (1:1000, Santa Cruz). Labeling by both primary antibodies was visualized using goat-anti-rabbit CY5 secondary antibody (1:500, Jackson ImmunoResearch Laboratories Inc., West Grove, PA, USA).

### Cell count analysis

Cells in individual taste buds were analyzed using Zen 2012 Blue Edition with the analysis done blind to experimental condition by 3 people. Images were taken at every micron and the nuclei in every 5^th^ slice was counted as an estimate of the number of taste cells within each bud. Cells expressing the target protein were then counted at every 5^th^ slice. Data is reported as the ratio of protein-expressing cells to total cells in the buds. Taste buds from at least 3 mice were analyzed for each antibody and experimental condition. Taste buds from 3 mice were also fixed, sectioned, labeled with DAPI and subsequently counted to compare the number of taste cells in the mice on the HF diet to mice on standard chow.

### cDNA synthesis

Isolated taste cells (see above) were placed in a microfuge tube and spun for 20 min at 13,000RPM to pellet cells. RNA was isolated using the Nucleospin RNA XS kit (Macherey-Nagel, Düren, Germany). RNA was treated with DNAase (Fermentas Life Sciences) and then used as a template to produce cDNA using SuperScript III Reverse Transcriptase (Invitrogen). PCR analysis was performed for GAPDH to ensure sample quality and check for genomic contamination. Contaminated samples were discarded and new samples were collected. Gustducin primers were taken from [42] and are listed in Supplemental Table 2. PLCβ2 and GAPDH primers are also listed in Supplemental Table 2.

### Real-Time PCR

Real-Time PCR was performed using a BioRad MiniOpticon system (Bio-Rad Laboratories, Hercules, CA), with BioRad SYBR Green reagents (Bio-Rad Laboratories). Taste cells from three mice were used for each sample and each sample was run in triplicate. If there was more than 5% difference between the replicates, the data were discarded. Data was normalized to GAPDH expression for each sample to correct for any loading differences and reported as fold differences. Three biological repeats were collected for each condition.

### Solutions

All chemicals were purchased from Sigma Aldrich (St. Louis, MO, USA) unless otherwise noted. Tyrode’s solution contained: 140mM NaCl, 5mM KCL, 1mM MgCl_2_, 3mM CaCl_2_, 10mM HEPES, 10mM glucose and 1mM pyruvic acid; pH 7.4. The following chemicals are diluted into Tyrode’s and used for cellular stimulation during experiments: Sweet-2mM saccharin (sac), 50mM sucrose (suc-50mM NaCl was replaced with 50mM sucrose) and 20mM acesulfame potassium (Ace K); Bitter-5mM denatonium benzoate (den). High potassium (Hi K) Tyrode’s (50mM NaCl replaced with 50mM KCL) was used to depolarize taste cells. The Ca^2+^-Mg^2+^-free solution consisted of 140mM NaCl, 5mM KCL, 10mM Hepes, 2mM EGTA, 2mM BAPTA, pH 7.4.

## Acknowledgments

The authors wish to thank Kristen E. Kay for her technical assistance. This work was supported by NSF funding 1256950 to Kathryn Medler, NIH DC006358 to Stefan Roberts and Kathryn Medler and DC016869 to A-M Torregrossa. We thank Alan Siegel and the UB North Campus Imaging Facility funded by NSF-MRI Grant DBI 0923133 for the confocal images.

## FIGURE LEGENDS

**Supplemental Figure 1:**
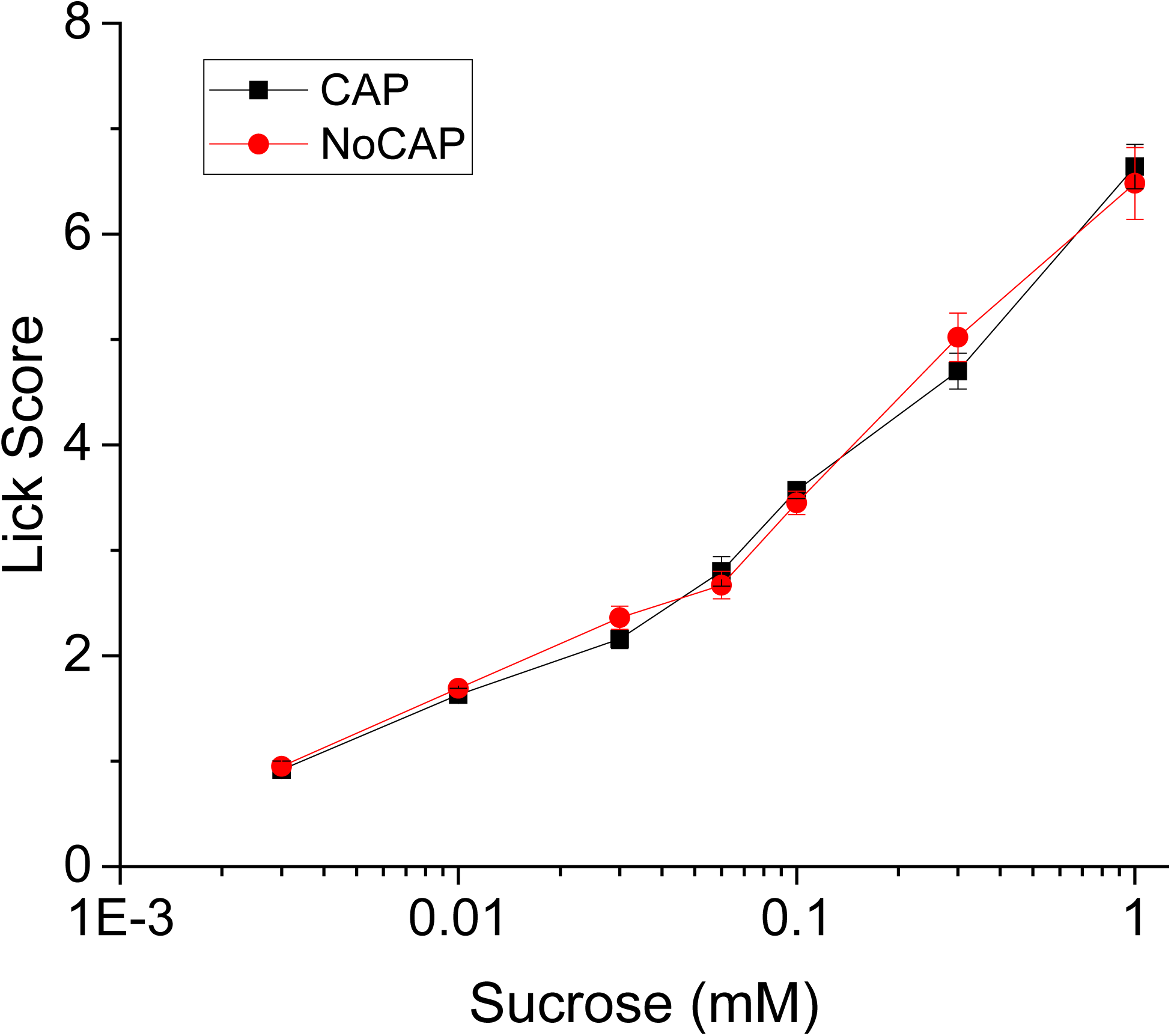
Captopril does not alter taste driven behaviors. Pre-exposure to CAP (4 days, 0.05mg/mL water) did not affect the unconditioned licking behavior of lean mice to sucrose (n=6 mice/group).

**Supplemental Table 1.** Raw data for taste cell responsiveness. The total number of cells tested and responses for each stimulus for each experimental condition is shown. Amplitude responses in Figures 2 and 3 were calculated from the responding cells in each condition and stimulus.

| Exp cond | AceK test | AceK resp | Sac test | Sac resp | Suc test | Suc resp | Den test | Den resp | KCl test | KCl resp |
| --- | --- | --- | --- | --- | --- | --- | --- | --- | --- | --- |
| HF | 150 | 10 | 152 | 27 | 169 | 13 | 172 | 23 | 166 | 16 |
| CTL | 80 | 11 | 72 | 25 | 80 | 10 | 80 | 22 | 80 | 12 |
| HF+CAP | 108 | 13 | 94 | 19 | 100 | 12 | 108 | 19 | 108 | 9 |
| CTL+CAP | 72 | 7 | 84 | 26 | 74 | 7 | 86 | 37 | 86 | 16 |

**Supplemental Table 2.**
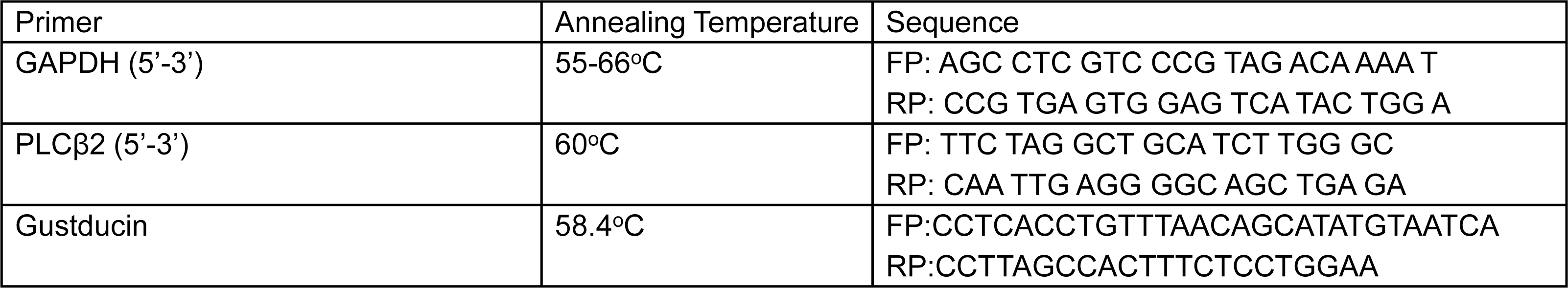
Real time primers sequences.

